# Distinct roles of the orbitofrontal cortex, ventral striatum, and dopamine neurons in counterfactual thinking of decision outcomes

**DOI:** 10.1101/2023.03.05.531219

**Authors:** Mengxi Yun, Masafumi Nejime, Takashi Kawai, Jun Kunimatsu, Hiroshi Yamada, HyungGoo R. Kim, Masayuki Matsumoto

## Abstract

Individuals often assess past decisions by comparing what was gained with what would have been gained had they acted differently. Thoughts of past alternatives that counter what actually happened are called “counterfactuals”. Recent theories emphasize the role of the prefrontal cortex in processing counterfactual outcomes in decision-making, although how subcortical regions contribute to this process remains to be elucidated. Here we report a clear distinction among the roles of the orbitofrontal cortex, ventral striatum and midbrain dopamine neurons in processing counterfactual outcomes in monkeys. Our findings suggest that actually-gained and counterfactual outcome signals are both processed in the cortico-subcortical network constituted by these regions but in distinct manners, and integrated only in the orbitofrontal cortex in a way to compare these outcomes. This study extends the prefrontal theory of counterfactual thinking and provides key insights regarding how the prefrontal cortex cooperates with subcortical regions to make decisions using counterfactual information.

**Teaser:** Cortical and subcortical systems both contribute to counterfactual thinking of decision outcomes but in distinct manners.

## Introduction

“If only I hadn’t sold my stock, I could have earned more money.” People often imagine how things could have turned out differently if they had made a different decision. Thoughts of what might have been, that is, thoughts of past alternatives that counter what actually happened, are called “counterfactuals” (*1*). Counterfactual thinking is an essential cognitive process in the assessment of past decisions (*1–3*). Rather than solely assessing past decisions based on the actual outcome (“what has been gained”), individuals can compare the outcome with counterfactual outcomes (“what might have been gained”) to reduce trial-and-error-based learning costs (*4*). Moreover, this type of comparison can trigger emotional experiences, such as regret, relief, or satisfaction, which can potentially have modulatory effects such as reinforcing good decisions and suppressing bad decisions in the future (*2, 5*).

How does the brain process counterfactual outcomes? The prefrontal cortex (PFC) is considered to be a promising candidate involved in this process. Indeed, functional magnetic resonance imaging (fMRI) studies in humans have revealed that multiple regions in the PFC are activated while subjects simulate counterfactual outcomes (*6–11*). More precise evidence from fMRI and single-unit recording studies in non-human primates has shown that neuronal activities in these prefrontal regions, including the orbitofrontal cortex (OFC) (*12*), anterior cingulate cortex (ACC) (*13, 14*), and dorsolateral prefrontal cortex (dlPFC) (*12*), represent the value of counterfactual outcomes. Consequently, it has been proposed that counterfactual outcomes are primarily processed by prefrontal networks.

A small number of studies have reported neural correlates of counterfactual information in subcortical regions. A previous study in human patients with Parkinson’s disease reported that subsecond dopamine (DA) release in the striatum not only represented conventional reward prediction error, i.e., the discrepancy between actual and expected reward values, but also reflected the value of counterfactual outcomes (*15*). Another study in rodents reported that neurons in the ventral striatum (VS) represented counterfactual choices that would have led to a better outcome than the actual outcome (*16*). However, how these subcortical signals are related to counterfactual outcome processing in the PFC is unclear. Notably, these subcortical regions are connected with prefrontal regions processing counterfactual outcomes, which together form a cortico-subcortical network that is essential for the expectation and evaluation of actually obtained rewards (*17, 18*). Therefore, it seems possible that counterfactual outcomes are processed by the homologous cortico-subcortical network governing actual reward computations. To investigate this possibility, it is necessary to understand how individual neurons in each component of this network represent actual and counterfactual outcomes.

In the present study, we aimed to address the above issues. Accordingly, we focused on the OFC, VS, and midbrain DA neurons that are major cortical and subcortical components of the cortico-subcortical reward network (*19–25*). We developed an economic decision-making task in which monkeys decided to choose or not to choose an option based on its value. After making a decision, a counterfactual or actual outcome was displayed. We compared the representations of the values of actual and counterfactual outcomes across the three cortical and subcortical regions, and found that the properties of their signals exhibited a clear contrast. While neurons in all three regions robustly represented the value of the actual outcome, they exhibited a descending cortico-subcortical gradient from the OFC, VS, to DA neurons in terms of the capacity to represent the value of the counterfactual outcome. Whereas the actual value signal appeared earliest in DA neurons, the counterfactual value signal appeared earliest in the OFC, suggesting different directions of information flow within the cortico-subcortical reward network between the actual and counterfactual value signals. Regarding the integration of actual and counterfactual outcome values, we found that the OFC, but not the other regions, represented the actual and counterfactual values in an antagonistic manner. This suggests that the OFC performs the comparison between these values, which is critical for the assessment of past decisions. In contrast, the VS represented the actual and counterfactual values in a common currency manner, as it individually evaluated these values. Our findings suggest that actual and counterfactual value processing are organized in parallel across the cortico-subcortical reward network and integrated in the OFC, which appears to perform critical comparison computations that support the assessment of past decisions.

## Results

### Effects of actual and counterfactual outcomes on monkey decision-making

We developed an economic decision-making task in which monkeys decided whether to choose an offered option. After they made a decision, a counterfactual outcome was displayed (Fig. 1A) (*26*). The monkey gazed at a central fixation point and pressed a button at the beginning of each trial, after which two of six possible visual objects were sequentially presented as first and second options. The six visual objects were associated with different amounts of a liquid reward. When the first option was presented, the monkey was required to decide whether to choose this option. Releasing the button during the presentation of the first option was regarded as the decision to choose the first option, while keeping the button pressed down was regarded as the decision not to choose this option. Then, the second option was presented. If the monkey had chosen the first option, it was unable to choose the second option even if the second option was better than the first option. If the monkey had not chosen the first option, it was required to release the button during the presentation of the second option, and this second option was regarded as being chosen. The monkey obtained the reward associated with the chosen option at the end of each trial.

**Fig. 1.**
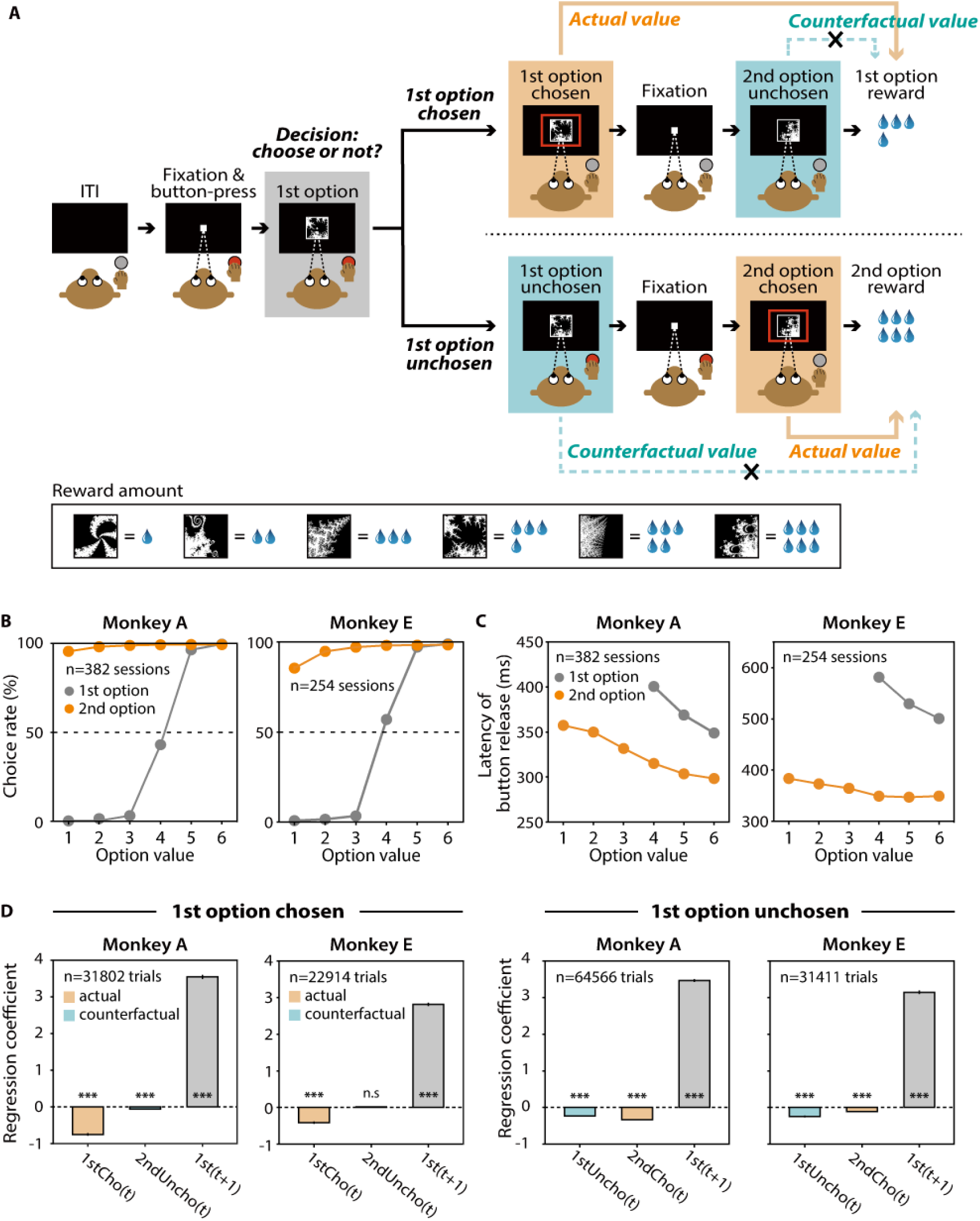
Effects of actual and counterfactual outcomes on monkey decisions. (A), Economic decision-making task. ITI, intertrial interval. First option chosen and first option unchosen trials are shown separately. (B), Choice rate of the first option (gray line) and choice rate of the second option (rate at which the monkey correctly released the button during the presentation of the second option in the first option unchosen trials) (orange line) in monkey A (left) and monkey E (right). (C), Latency of button release to choose the first (gray line) and second (orange line) option in monkey A (*N* = 381 sessions) (left) and monkey E (*N* = 255 sessions) (right). (D), Effects of the actual outcome value (first and second option values in the first option chosen and unchosen trials, respectively) (orange) and the counterfactual value (second and first option values in the first option chosen and unchosen trials, respectively) (green) in trial t and the first option value in trial t+1 (gray) on the monkey’s decision in trial t+1. These effects were calculated separately for the first option chosen (left) and unchosen (right) trials for each monkey. Triple asterisks indicate a significant logistic regression coefficient (*P* < 0.001; t-test). n.s. indicates no significance (*P* > 0.05; t-test). Error bars indicate SEM, which were very small and hidden in most cases.

In trials in which the monkey chose the first option (hereafter called “first option chosen trials”), the outcome associated with the first option became “actual” when the monkey decided to choose it, while the outcome associated with the second option was “counterfactual”. In trials in which the monkey did not choose the first option (hereafter called “first option unchosen trials”), the outcome associated with the first option became “counterfactual” when the monkey decided not to choose it, while the outcome associated with the second option was “actual”. Thus, the associations between the actual/counterfactual outcomes and the first/second options were determined when the monkey finalized its decision.

We trained two monkeys (monkeys A and E) to perform this decision-making task. When the first option had a higher value, the monkeys were more likely to choose this option (logistic regression slope; monkey A, means ± standard deviation [SD] = 3.8 ± 0.9, *P* < 0.0001; monkey E, means ± SD = 3.4 ± 1.0, *P* < 0.0001; two-tailed Wilcoxon signed-rank test) (gray line in Fig. 1B). In addition, the latency of the action performed to choose the first option (i.e., button release) decreased as the value of the first option increased (regression slope; monkey A, means ± SD = −25.3 ± 8.7, *P* < 0.0001; monkey E, means ± SD = −39.1 ± 23.8, *P* < 0.0001; two-tailed Wilcoxon signed-rank test) (gray line in Fig. 1C). If the monkey did not choose the first option, it needed to release the button during the presentation of the second option to choose the second option. However, the monkeys sporadically failed to release the button, especially when the value of the second option was low (regression slope; monkey A, means ± SD = 0.7 ± 0.9, *P* < 0.0001; monkey E, means ± SD = 2.2 ± 2.9, *P* < 0.0001; two-tailed Wilcoxon signed-rank test) (orange lines in Fig. 1B and fig. S1A). Because decision-making was not required when the monkey released the button to select the second option, the button release latency for the second option was shorter than that for the first option (monkey A, value 4: *P* < 0.0001, value 5: *P* < 0.0001, value 6: *P* < 0.0001; monkey E, value 4: *P* < 0.0001, value 5: *P* < 0.0001, value 6: *P* < 0.0001; two-tailed Wilcoxon signed-rank test)) (Fig. 1C). The button release latency for the second option decreased as the value of the second option increased (regression slope; monkey A, means ± SD = −13.1 ± 5.8, *P* < 0.0001; monkey E, means ± SD = −7.3 ± 7.3, *P* < 0.0001; two-tailed Wilcoxon signed-rank test) (orange lines in Fig. 1C). These data indicate that the monkey choice behavior was affected by the value of the chosen option, which was the actual outcome value.

How did the counterfactual outcome influence the monkey behavior during the task? Usually, counterfactual outcomes influence future decisions (*7, 14, 27*). To address this, we examined the effect of the counterfactual outcome on monkey decision-making in the next trial. We examined this effect separately for the first option chosen trials and first option unchosen trials because the counterfactual outcome was associated with different options (the second and first options, respectively) (Fig. 1A). For each trial type, we used a logistic regression model that predicted whether the monkey chose the first option in trial t+1 based on (1) the value of the first option in trial t+1, (2) the value of the first option in trial t, and (3) the value of the second option in trial t (Materials and Methods). The values of the first and second options in trial t are the values of the actual and counterfactual outcomes, respectively, in the first option chosen trials (and vice versa in the first option unchosen trials). We observed that the decision in trial t+1 was most strongly affected by the value of the first option in trial t+1 in both trial types (regression coefficient for the first option chosen trials; monkey A, means ± standard error of the mean [SEM] = 3.5 ± 0.05, *P* < 0.0001; monkey E, means ± SEM = 2.8 ± 0.04, *P* < 0.0001; regression coefficient for the first option unchosen trials; monkey A, means ± SEM = 3.5 ± 0.04, *P* < 0.0001; monkey E, means ± SEM = 3.1 ± 0.04, *P* < 0.0001; t test) (gray bars in Fig. 1D), indicating that whether the monkey chose the first option was mainly determined by the value of the first option. However, the values of the actual and counterfactual outcomes in the last trial also influenced the monkey’s decision. That is, the effect of the actual outcome value in trial t was significant in both trial types (regression coefficient for the first option chosen trials; monkey A, means ± SEM = −0.75 ± 0.03, *P* < 0.0001; monkey E, means ± SEM = −0.04 ± 0.03, *P* < 0.0001; regression coefficient for the first option unchosen trials; monkey A, means ± SEM = −0.34 ± 0.01, *P* < 0.0001; monkey E, means ± SEM = −0.1 ± 0.01, *P* < 0.0001; t test) (orange bars in Fig. 1D). Notably, the effect of the actual outcome value in trial t was opposite that of the first option value in trial t+1. This indicates that as the value of the actual outcome in the last trial increased, the chance that the monkey would choose the first option in the next trial decreased.

The effect of the counterfactual value in trial t was different between the first option chosen and the first option unchosen trials. In the first option unchosen trials, the effect of the counterfactual outcome value (i.e., the value of the first option) was significant (regression coefficient; monkey A, means ± SEM = −0.23 ± 0.018, *P* < 0.0001; monkey E, means ± SEM = −0.25 ± 0.018, *P* < 0.0001; t test) (green bars in Fig. 1D). This effect was opposite that for the value of the first option in trial t+1, as seen in the actual outcome values (orange bars in Fig. 1D). This indicates that as the value of outcome that the monkey saw as counterfactual increased, the chance that the monkey would choose the first option in the next trial decreased. However, in the first option chosen trials, the effect of the counterfactual outcome value (i.e., the value of the second option) was weak, and it was significant in only one of the two monkeys (regression coefficient; monkey A, means ± SEM = −0.060 ± 0.015, *P* < 0.0001; monkey E, means ± SEM = 0.018 ± 0.016, *P* = 0.26; t test) (green bars in Fig. 1D). The difference between these trial types could be explained by the “behavioral demand” of the option associated with the counterfactual outcome. In the first option unchosen trials, the counterfactual outcome was associated with the first option, and the monkey needed to evaluate this option to decide not to choose it. In contrast, in the first option chosen trials, the counterfactual outcome was associated with the second option, and this option was presented after the monkey had decided to choose the first option (i.e., after the monkey had decided not to choose the second option). Thus, the monkey did not need to evaluate the second option, at least for the decision-making in the ongoing trial. That the behavioral demand of the second option was lower than that of the first option is one possible explanation for the weaker effect of the counterfactual value of the second option on the monkey’s decision in the next trial.

### OFC, VS, and DA neurons robustly represented the value of the actual outcome

We recorded single unit activity from 96 DA neurons (60 and 36 neurons in monkeys A and E, respectively) in the substantia nigra pars compacta (SNc) and ventral tegmental area (VTA), 285 neurons in the OFC (156 and 129 neurons in monkeys A and E), and 255 neurons in the VS (166 and 89 in monkeys A and E) (fig. S2) (Materials and Methods). We found that many of these neurons showed activity that was modulated by the values of the first and/or second option, although the modulation pattern was somewhat diverse depending on the monkey’s decision (Fig. 2). For instance, an example DA neuron increased its activity as the value of the first option increased during the presentation of the first option in both the first option chosen and unchosen trials, indicating that this neuron represented the value (left column in Fig. 2A). However, this does not guarantee that this DA neuron represented the counterfactual value of the first option, because the first option had not yet been counterfactual before the monkey decided not to choose this option. Indeed, this DA neuron did not represent the counterfactual value of the second option, which was counterfactual from its onset in the first option chosen trials (right column in Fig. 2A). An example VS neuron represented the values of the first and second options when the monkey chose these options, although this neuron did not represent the values when the monkey did not choose the first option (left column in Fig. 2B) or when the second option was counterfactual (right column in Fig. 2B). Contrary to the activity of these neurons, some of the recorded neurons represented the counterfactual value of the second option. An example OFC neuron increased its activity as not only the actual value but also the counterfactual value increased, indicating that this neuron represented these values in the same fashion (right column in Fig. 2C).

**Fig. 2.**
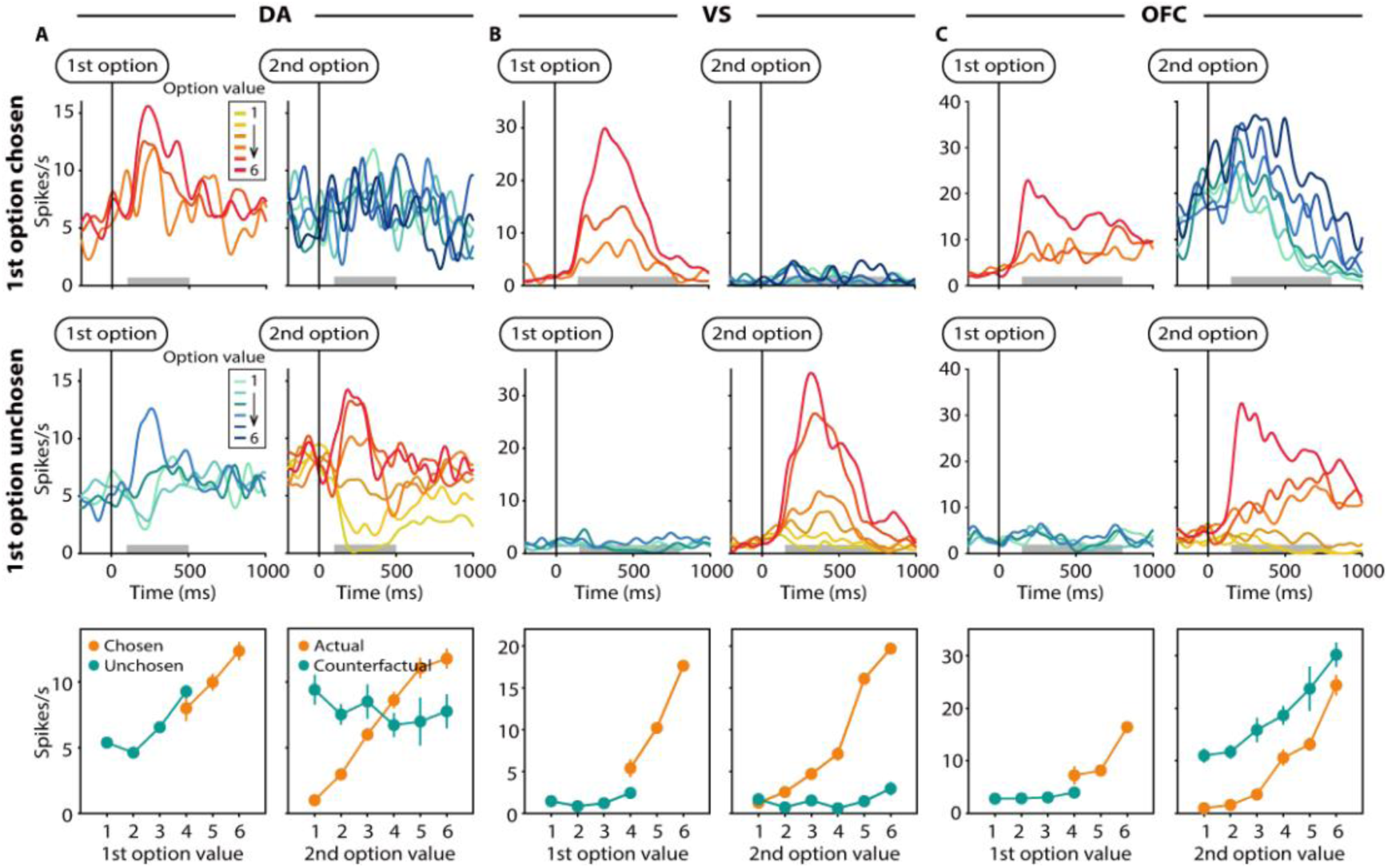
OFC, VS, and DA neurons representing the chosen/unchosen value of the first option and the actual/counterfactual value of the second option. (A to C), Top and Middle: Activity of example neurons (A, DA neuron; B, VS neuron; C, OFC neuron) in the first option chosen trials and first option unchosen trials, respectively. Spike density functions (SDFs) are aligned at the onsets of the first (left) and second (right) options. The orange and blue SDFs indicate the activity when the first option was chosen and not chosen, respectively, and when the second option was actual and counterfactual, respectively. The darker the color, the higher the value of each option. Gray horizontal bars indicate the time window used to calculate the magnitude of neuronal activity for each value. Bottom: Magnitudes of neuronal activity plotted against the value for the chosen (orange) and unchosen (blue) first options and for the actual (orange) and counterfactual (blue) second options. Error bars indicate SEM.

To statistically identify the value representation of counterfactual outcomes in the three brain regions, we focused on neuronal modulations evoked by the second option, despite its weak effect on the next decision, since neuronal modulations evoked by the first option did not purely reflect counterfactual outcome information. Although the first option became counterfactual in the first option unchosen trials, this option was not counterfactual before the monkey decided not to choose it. Besides, it was not possible to measure the time point at which the monkey decided to not to choose this option, because the monkey expressed this decision by keeping the button pressed down (i.e., no action was required for this decision). In the first option chosen trials, in contrast, the second option was counterfactual from its onset. Thus, we used the neuronal modulation evoked by the second option to investigate the neural mechanisms underlying counterfactual value processing. We used a calculation time window set after the onset of the second option, during which neuronal modulations evoked by the second option were observed as a population, and we fit the activity of each neuron during this window using a generalized linear model (GLM) separately for the two decision trial types (Materials and Methods). In the GLM, the values of the first and second options were set as regressors. If the weight of the second option value in the first option chosen and unchosen trials was significant in the fitted model, that neuron was considered to represent the actual and counterfactual values of the second option, respectively.

As expected, we first identified a substantial proportion of neurons representing the actual value of the second option in the OFC, VS, and DA neurons (OFC, 47.47%, *P* < 0.001; VS, 53.27%, *P* < 0.001; DA, 66.67%, *P* < 0.001; one-tailed bootstrap test) (Materials and Methods) (Fig. 3A). Among these regions, the actual value representation in most of the DA neurons had a positive weight (positive weight: 92.19%, negative weight: 7.81%), whereas the VS and OFC contained similar proportions of neurons with positive and negative weights (VS, positive weight: 53.51%, negative weight: 46.49%; OFC, positive weight: 47.54%, negative weight: 52.46%) (Fig. 3A). Fig. 3B shows the spike density functions (SDFs) of neurons in the three regions that positively represented the actual value of the second option (see also top in fig. S3 for neurons that negatively represented the value) (Materials and Methods). These results are consistent with a rich body of work that has demonstrated the crucial involvement of the OFC, VS, and DA neurons in actual reward computations.

**Fig. 3.**
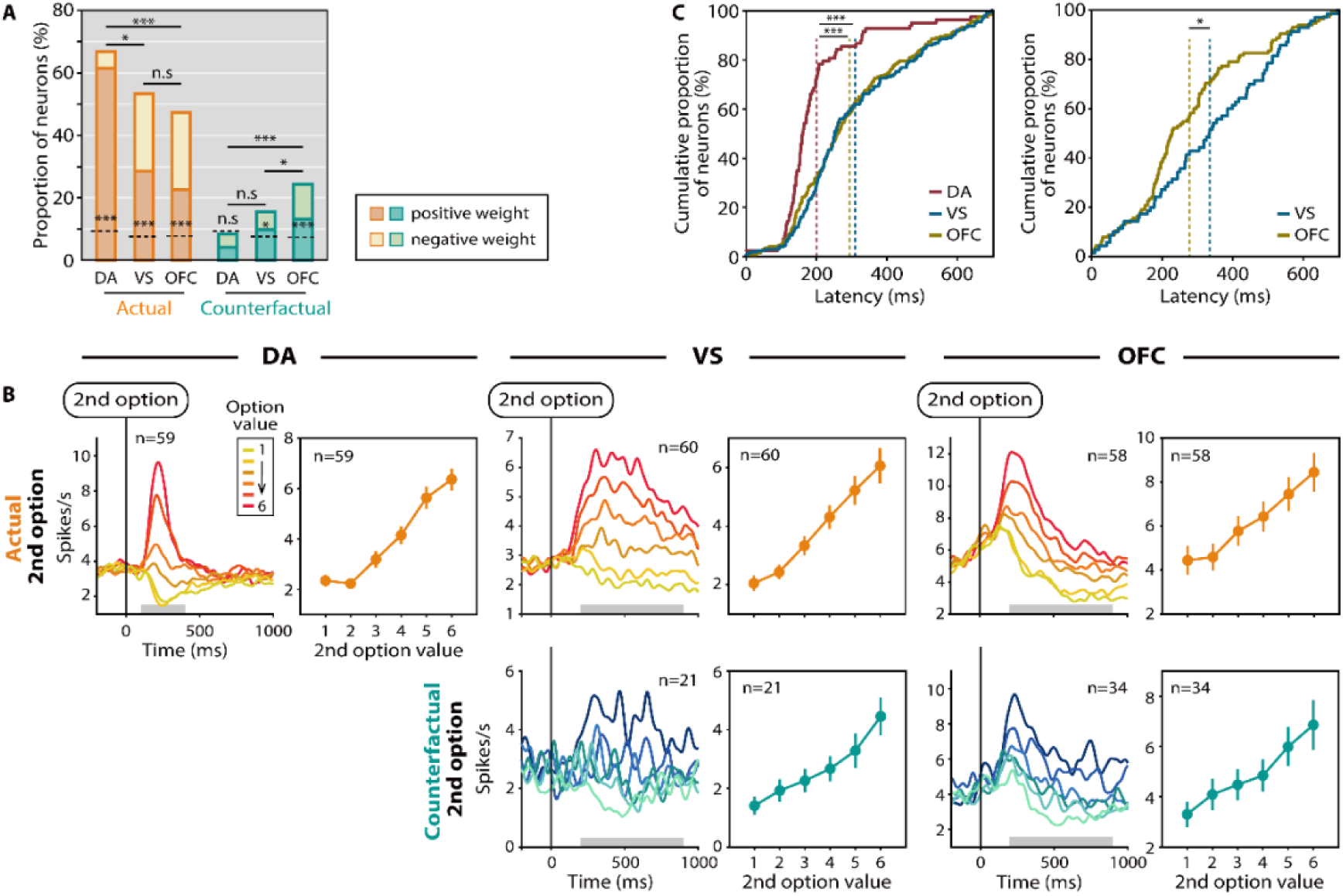
Representation of actual and counterfactual second option values in OFC, VS, and DA neurons. (A), Proportions of OFC, VS, and DA neurons representing the actual (orange) and counterfactual (blue) values of the second option. Bars with dark and light colors indicate neurons for which activity increased and decreased, respectively, as the value increased (positive and negative types, respectively). Dotted lines indicate the chance level calculated by a bootstrap procedure. Single and triple asterisks above the dotted lines indicate a significantly larger proportion than the chance level (*P* < 0.05 and 0.001, respectively; one-tailed bootstrap test). Single and triple asterisks between bars indicate a significant difference between the proportions (*P* < 0.05 and 0.001, respectively; two-tailed Fisher’s exact test). n.s. indicates no significance. (B), Averaged SDFs aligned at the second option onset (left) and average magnitudes of neuronal activity evoked by the second option (right) for the first option unchosen and chosen trials in which the second option was the actual (top) and counterfactual (bottom) outcome, respectively. Neurons that positively represented the second option value were used in this analysis (see fig. S3 for neurons that negatively represented the value). Conventions are as Fig. 2. (C), Cumulative distributions of the latencies of the actual (left) and counterfactual (right) value signals shown for OFC (yellow line), VS (blue line), and DA neurons (red line) (actual: OFC, *N* = 122, VS, *N* = 114, DA, *N* = 64; counterfactual: OFC, *N* = 63, VS, *N* = 33). Vertical dotted lines indicate mean latencies. Single and triple asterisks indicate a significant difference between the latencies (*P* < 0.05 and 0.001, respectively; two-tailed Wilcoxon signed-rank test).

### OFC, VS, and DA neurons exhibited a descending gradient in the capacity to represent the value of the counterfactual outcome

In addition to the value of the actual outcome, we also examined whether representations of the value of the counterfactual outcome are shared across these regions. Analyzing the GLM coefficients, we found that significant proportions of OFC neurons represented the counterfactual value of the second option, even if the monkey could only see but not choose the option (24.51%, *P* < 0.001; one-tailed bootstrap test) (Fig. 3A) (see right bottom in Fig. 3B for SDFs of OFC neurons that positively represented the value, bottom in fig. S3B for OFC neurons that negatively represented the value). Moreover, not only in the OFC but also in the subcortical VS, we found a significant proportion of neurons that represented the counterfactual value of the second option (15.42%, *P* = 0.015; one-tailed bootstrap test) (Fig. 3A) (see middle bottom in Fig. 3B for SDFs of VS neurons that positively represented the value, bottom in fig. S3A for VS neurons that negatively represented the value), although the proportion of neurons that represented the counterfactual value of the second option was significantly lower in the VS than in the OFC (*P* = 0.016, Fisher’s exact test). In contrast to these two regions, the proportion of DA neurons representing the counterfactual value of the second option was not statistically different from the chance level calculated using a bootstrap procedure (8.33%, *P* = 1.0; one-tailed bootstrap test) (Fig. 3A). This indicates that DA neurons represented the value of the outcome only when the outcome was actually available. Our findings suggest a descending gradient in the capacity to represent the value of the counterfactual outcome from the OFC, VS, to DA neurons.

As mentioned, the effect of the counterfactual value of the second option on the next decision was weak and significant in only one of the two monkeys (Fig. 1D) (see also fig. S4 for the proportion of neurons representing the counterfactual value for each monkey, which was not significantly different between the two monkeys). As discussed, this weak effect may be explained by the low behavioral demand of this option. Nevertheless, we found that significant numbers of OFC and VS neurons represented the counterfactual value. Thus, these regions might evaluate the counterfactual value regardless of the degree of behavioral demand.

We compared the latencies of the actual and counterfactual value signals across the three regions (Fig. 3C). The latency of the actual value signal was significantly shorter in DA neurons (means ± SEM = 200.24 ± 127.43 ms) than in the OFC (means ± SEM = 293.94 ± 164.62 ms) and the VS (means ± SEM = 310.60 ± 168.93 ms) (DA versus OFC, *P* < 0.0001; DA versus VS, *P* < 0.0001; OFC versus VS, *P* = 0.40; two-tailed Wilcoxon signed-rank test). By contrast, the latency of the counterfactual value signal was significantly shorter in the OFC (means ± SEM = 276.46 ± 172.31 ms) compared with the latency in the VS (means ± SEM = 335.27 ± 182.98 ms) (*P* = 0.023; two-tailed Wilcoxon signed-rank test). This suggests that actual and counterfactual value signal transmissions have different directions of information flow between the cortical and subcortical regions.

### Neurons in the OFC and VS represented actual and counterfactual values in a common currency-like format

Given our finding that neurons in the OFC and VS represented both the actual and counterfactual values of the second option, we next examined how individual neurons in these regions organized the two types of values. For instance, do the same neurons represent both values? Or, do different populations of neurons represented each value? As mentioned above, we calculated the regression coefficients between neuronal activity and actual/counterfactual values (i.e., the weights of the regressors in the GLM) for each neuron. We then plotted these coefficients as scatterplots for each region (Fig. 4). In DA neurons, the plots expanded along the right side of the x-axis, indicating that the activity of these neurons increased as the actual value increased but did not change as a population in response to the counterfactual value. There was no significant correlation between the coefficients (*P* = 0.47; left in Fig. 4). However, we found a significantly positive correlation between the coefficients of the actual and counterfactual values in both the OFC and VS (VS: *P* < 0.0001, middle in Fig. 4; OFC: *P* < 0.0001, right in Fig. 4). That is, OFC and VS neurons of which activity changed with the actual value of the second option were more likely to exhibit an activity modulated with the counterfactual value of the second option with the same tuning direction. This correlation argues against the possibility that actual and counterfactual values are represented in different neuron populations in the OFC and VS. Instead, it suggests that OFC and VS neuron populations generalize actual and counterfactual values by representing both values in a common currency-like format.

**Fig. 4.**
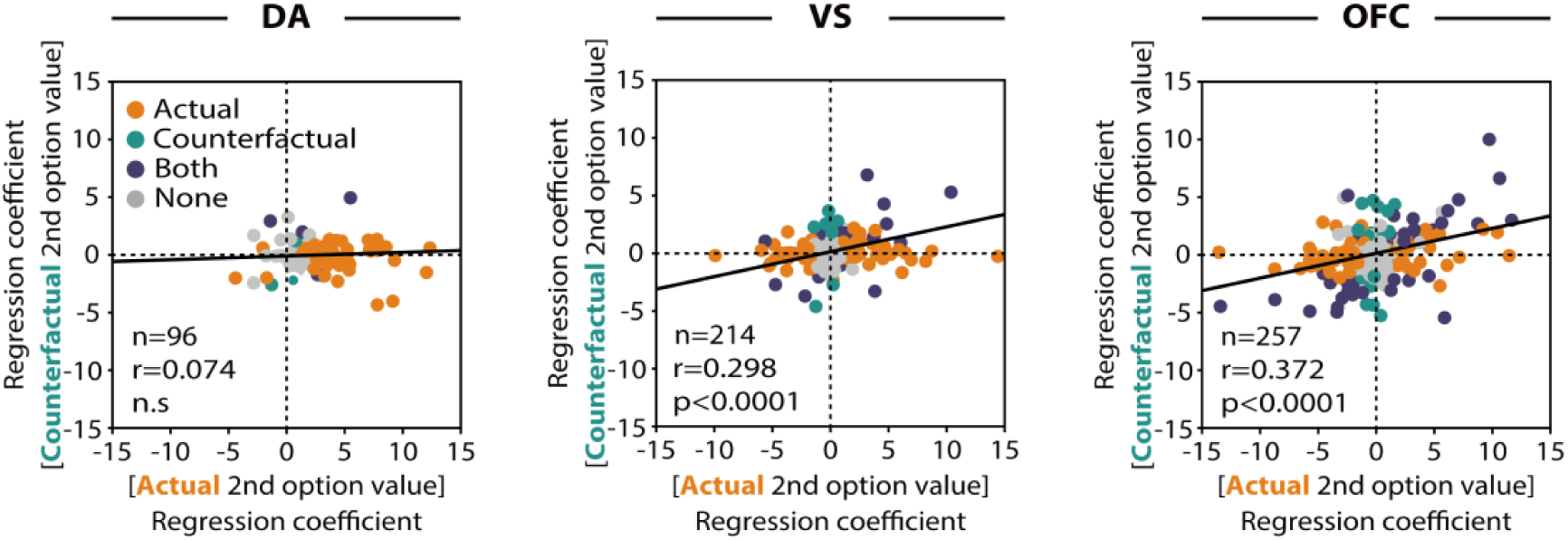
Relationship between actual and counterfactual value representations in OFC and VS neurons. Regression coefficients for the actual (x-axis) and counterfactual (y-axis) values of the second option are plotted. Each plot indicates each neuron. Orange, blue, purple, and gray plots indicate neurons with significant regression coefficient(s) for the actual value, counterfactual value, both values, and neither values, respectively (*P* < 0.05; t test). The regression line is shown as a black line.

### Counterfactual value representation in the OFC was influenced by the value of the actually chosen option in an antagonistic manner

Counterfactual thinking assesses past decisions and triggers emotional experiences such as regret by comparing what has been gained (actual outcome) with what might have been gained (counterfactual outcome) (*28*). We next investigated whether and how the OFC, VS, and DA neurons are involved in this comparison. Fig. 5A shows the neuronal dynamics related to signaling the value of the chosen first option (actual value) and the value of the unchosen second option (counterfactual value) in the first option chosen trials. Here, we used a GLM with these two values as regressors and analyzed the proportion of neurons showing a significant regression coefficient for each value with a sliding time window (Materials and Methods). Notably, the actual value signal in the OFC evoked by the first option remained even after the onset of the second option, at which point counterfactual processing could have begun, and coexisted with the counterfactual value signal evoked by the second option (right in Fig. 5A). To examine how these different types of value signals were integrated into neuronal activity, we plotted the regression coefficient of the actual value (x-axis) and that of the counterfactual value (y-axis) as scatterplots for each neuron (Fig. 5B). We found a significantly negative correlation between the coefficients in the OFC but not in the VS or DA neurons (DA: *P* = 0.80; VS: *P* = 0.35; OFC: *P* < 0.0001), indicating that OFC neurons represented the actual and counterfactual values in an antagonistic manner. Such antagonistic representation of two different values has been considered as a potential neural mechanism for comparing the values (*29–31*). Therefore, the observation of antagonistic representation in the OFC raises the possibility that this prefrontal region is involved in the comparison of the actual and counterfactual values, which is a key process involved in counterfactual thinking.

**Fig. 5.**
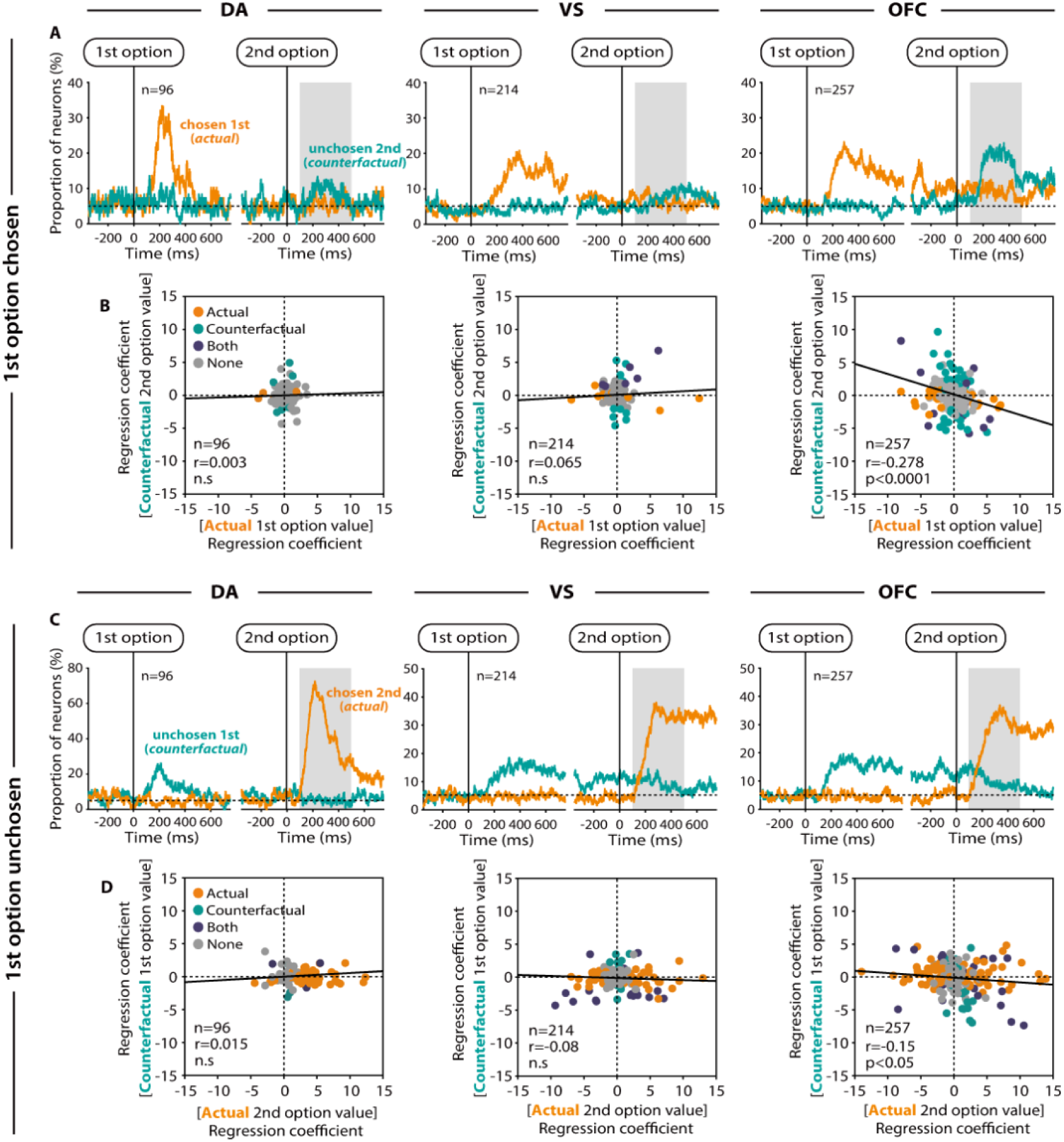
Antagonistic representation of the actual and counterfactual values of competing options in OFC neurons. (A and C), Temporal change in the proportions of neurons representing the first and second option values, aligned at the first and second option onsets. Proportions are shown for the first option chosen (A) and unchosen (C) trials. Gray areas indicate the time windows used to calculate the scatterplots shown in (B) and (D). Horizontal dotted lines indicate the baseline proportion of neurons representing the value of options, calculated using a time window prior to the first option onset. (B and D), Scatterplots of the regression coefficients of the actual (x-axis) and counterfactual (y-axis) values in the first option chosen (B) and unchosen (D) trials. Conventions are as Fig. 4.

We next examined the neuronal dynamics signaling the actual and counterfactual values in the first option unchosen trials, in which the actual outcome was associated with the second option and the counterfactual outcome was associated with the first option (Fig. 5C). Even in these trials, the counterfactual value signal in the OFC evoked by the first option remained even after the onset of the second option (right in Fig. 5C). We found a weak negative correlation between the coefficients of the actual and counterfactual values in the OFC (DA: *P* = 0.26; VS: *P* = 0.11; OFC: *P* = 0.016) (right in Fig. 5D). Taken together, our findings suggest that while the representation of actual and counterfactual values appears to be shared across the OFC and subcortical VS, the process of comparing these values may be primarily localized in the OFC.

## Discussion

In the present study, we investigated the role of key nodes in the cortico-subcortical reward network, i.e., the OFC, VS, and DA neurons, in processing the value of counterfactual outcomes in economic decision-making. We found a clear contrast among their roles in processing the counterfactual value. Although all of these cortical and subcortical regions robustly represented the value of actual outcomes, their capacity to represent the counterfactual value showed a descending cortico-subcortical gradient from the OFC, VS, to DA neurons (Fig. 3A). While the actual value signal appeared earliest in DA neurons, the counterfactual value signal appeared earliest in the OFC (Fig. 3C). Moreover, neurons in the VS and OFC represented the actual and counterfactual values in a unified manner (Fig. 4), although the neural signature of comparison between these values, which is a critical process in the assessment of past economic decision-making, was only observed in the OFC (Fig. 5). The different features observed for counterfactual value processing among the OFC, VS, and DA neurons were distinct in terms of their robust representation of actual value. This provides important insight regarding how the prefrontal cortex, which has singularly attracted attention as the neural substrate of counterfactual thinking, cooperates with other brain structures to make better decisions based on counterfactual information about past decision outcomes.

A previous human imaging study (*6*) and a non-human primate electrophysiological study (*12*) have reported that the OFC is involved in the processing of both actual and counterfactual outcomes. However, the computations executed by neurons in the OFC to process and integrate actual and counterfactual values are still unclear. In the present study, we compared the ways in which OFC neurons represented actual and counterfactual values associated with the same option (i.e., second option) in different contexts, and found that these neurons represented the two values in a unified manner (Fig. 4). This suggests that they use a common-currency format to generate the representation of the two value types. We found that individual OFC neurons represented the actual and counterfactual values in an antagonistic manner if these values were associated with different options (i.e., first and second options) that were competing in a trial (Fig. 5). Such antagonistic representation of two different types of values is regarded as a neural signature of comparison between these values (*29–31*). Previous studies have reported that OFC neurons exhibited another antagonistic value representation of two offered options in economic decision-making tasks (*31, 32*). This antagonistic value representation occurred before the subject decided which option to choose, and is thought to play a key role in comparing two options during ongoing decision-making. Taken together with our finding that the antagonistic representation of the actual and counterfactual values occurred after the monkey had made a decision, these data indicate that the OFC broadly participates in comparison processes during economic decision-making.

In addition to those in the OFC, we found that neurons in the VS represented the actual and counterfactual values of the second option in a unified manner. The VS is well-known for its functions in reward prediction, reward-based learning, and action selection (*33, 34*). Anatomically, the VS is a major target of projections from medial prefrontal areas including the OFC (*17, 35–37*), in which a policy is formulated to maximize reward gain according to different external contextual and internal state information (*38, 39*). Given the existence of projections from the prefrontal areas to the VS, our findings regarding counterfactual value representation in the VS and a longer latency for the counterfactual value signal in the VS compared with the OFC are consistent with the anatomy. Therefore, counterfactual value signals may first be processed in prefrontal areas such as the OFC and then transmitted to the subcortical VS. However, the proportion of VS neurons representing the counterfactual value was significantly smaller than that for OFC neurons (Fig. 3A). Furthermore, although the OFC exhibited a neural signature of actual and counterfactual value comparison (i.e., the antagonistic representation of the actual and counterfactual values of competing options), this was not the case for the VS (Fig. 5B and D). Our findings regarding the reduced proportion and lack of integration between the actual and counterfactual values might reflect a main feature of the VS. That is, as previously proposed, the VS might trigger motivational actions such as reward seeking and punishment avoidance (*40, 41*). Counterfactual outcomes are not directly associated with triggering these motivational actions, but are instead useful for internal simulations that enable decisions about future actions.

We found that DA neurons only represented actual values and not counterfactual values in our economic decision-making task (Fig. 3A). DA neurons have been shown to encode reward prediction error, which indicates the discrepancy between actual and expected reward values, and are thought to update expectations about future rewards (*21*). Another type of prediction error using counterfactual reward information instead of actual reward information, i.e., the discrepancy between counterfactual and expected reward values, may also be useful in updating expectations about future rewards (*12, 27, 42*). However, our findings suggest that DA neurons do not encode this type of prediction error. Different brain regions might provide distinct prediction errors to enhance the accuracy of future reward expectations according to multiple aspects of economic decision-making. In terms of systems neuroscience focused on economic decision-making, this “negative” finding provides an important insight regarding the boundaries of DA signal function in reinforcement learning.

The effect of counterfactual outcomes on future decisions has been reported in multiple animal species, including humans (*5, 6*), non-human primates (*12, 33, 43*), and rats (*27*). Instead of solely using the values of actually chosen outcomes, previous studies have shown that adding the value of an unchosen option (i.e., counterfactual value) into models produced a better fit with respect to animal choice behavior. In the present study, we analyzed the effect of the counterfactual outcome value on monkey decisions in the next trial for “first option chosen trials” and “first option unchosen trials” separately because the counterfactual outcome was associated with different options (the second and first options, respectively) (Fig. 1D). The effect was particularly significant in the first option unchosen trials, while weak and significant in only one of the two monkeys in the first option chosen trials. As mentioned in the result section, the weak effect in the first option chosen trials could be explained by the low behavioral demand of the option associated with the counterfactual outcome. That is, in the first option chosen trials, the counterfactual outcome was associated with the second option, which was only presented after the monkey had decided to choose the first option (i.e., after the monkey had decided not to choose the second option). Thus, the monkey did not need to evaluate the second option as part of the ongoing decision-making. The low behavioral demand of the second option might have influenced the weak effect of the counterfactual value on the monkey’s decision in the next trial.

Despite its weak effect on the monkey’s next decision, we mainly focused on the neuronal modulation evoked by the second option for the following reason. In the first option unchosen trials, the first option had not yet been counterfactual before the monkey decided not to choose it. We could not measure the time at which the monkey made this decision because the decision not to choose the first option was expressed by keeping the button pressed down (i.e., no action was required). Thus, it is difficult to extract the effect of the counterfactual value associated with the first option on neuronal activity. In the first option chosen trials, however, the second option was counterfactual from its onset. Thus, we expected the neuronal modulation evoked by the second option to purely reflect the counterfactual value. We found that significant numbers of OFC and VS neurons represented the counterfactual value of the second option. Our findings suggest that neurons in the OFC and VS evaluate the counterfactual value regardless of the degree of its effect on future decisions. It should be noted, however, that the proportion of neurons, including DA neurons, representing counterfactual value may change during decision-making tasks in which the counterfactual value strongly influences animal behavior.

In summary, thinking of what we would have gained had we acted differently, i.e., counterfactual outcomes, requires abstract internal representation and retrospective recognition. The present study uncovered a clear distinction between the roles of cortical and subcortical elements of the cortico-subcortical reward network in counterfactual value processing. Our findings suggest that actual and counterfactual value signals are processed in distinct manners in the cortico-subcortical reward network, and are integrated in an antagonistic comparison format that is present only in the PFC. This study provides key insights regarding the circuit-level neural mechanism underlying counterfactual thinking during economic decision-making.

## Materials and Methods

### Database

In the present study, we used a database obtained in our previous study in which single-unit activity was recorded from the orbitofrontal cortex (OFC) and dopamine (DA) neurons in monkeys performing an economic decision-making task (*26*). We newly recorded single-unit activity from the ventral striatum (VS) in the same monkeys. The procedures of the behavioral task and electrophysiology were identical to those in the previous study (*26, 44*).

### Animals

Two adult rhesus monkeys (Macaca mulatta; monkey A, male, 8.6 kg, 6 years old; monkey E, male, 10.1 kg, 12 years old) were used for the experiments. All procedures for animal care and experimentation were approved by the University of Tsukuba Animal Experiment Committee (permission number, 14-137), and were carried out in accordance with the guidelines described in Guide for the Care and Use of Laboratory Animals published by the Institute for Laboratory Animal Research.

### Behavioral tasks

Behavioral tasks and data collection were controlled by TEMPO system (Reflective Computing, WA, USA). The monkeys sat in a primate chair facing a computer monitor in a sound-attenuated and electrically shielded room. Eye movements were monitored using an infrared eye-tracking system (Eyelink, SR research, Ontario, Canada) with sampling at 500 Hz.

The monkeys were trained to perform an economic decision-making task (Fig. 1A). Six visual objects were associated with different amounts of a liquid reward (0.12, 0.18, 0.24, 0.3, 0.36, and 0.42 ml). The visual objects were monochrome fractal images (width, 5.2 deg; height, 5.2 deg) in monkey E, and bar stimuli (width, 5.3 deg; height, 2.3 deg) consisting of green and magenta areas, the fraction of which predicted the amount of the liquid reward, in monkey A. Each trial began with the presentation of a central fixation point (diameter, 0.5 deg) on the monitor, and the monkey was required to fixate on the point and press a button with the right hand. After the monkey had maintained fixation and kept the button pressed down for 750 ms, the fixation point disappeared and one of the six visual objects was randomly presented as the “first option” at the center of the monitor for 1000 ms. The monkey was required to decide to choose or not to choose this first option within its presentation. Releasing the button was regarded as the decision to choose the first option, while keeping the button pressed down was regarded as the decision not to choose it. When the monkey released the button, a red open rectangle (width, 6.3 deg; height, 6.3 deg) was presented around the chosen option as feedback. The first option and red rectangle disappeared 1000 ms after the onset of the first option, followed by a 400-ms fixation period. Then, one of the six visual objects was randomly presented as the “second option” for 1000 ms. If the monkey had decided to choose the first option (“first option chosen trials”), the animal was required to maintain the fixation to the second option during its presentation but could not choose it regardless of the second option value and obtained the reward associated with the first option after the presentation of the second option. If the monkey had decided not to choose the first option (“first option unchosen trials”), the animal obtained the reward associated with the second option by releasing the button within the presentation of the second option. When the monkey released the button, the red open rectangle was presented around the second option as feedback. The monkey was required to maintain fixation until the offset of the second option. Correct behavior was signaled by a tone (1 kHz), and the reward associated with the chosen option was simultaneously delivered. Trials were aborted immediately if the monkey (1) did not start the central fixation or press the button within 4000 ms after the onset of the fixation point, (2) broke the central fixation, (3) released the button during inappropriate periods (i.e., during the first and second fixation periods), (4) pressed the button twice, or (5) failed to release the button during a trial. These errors were signaled by a beep tone (100 Hz). All trials were presented with a random intertrial interval (ITI) ranging from 2000 to 3000 ms.

### Electrophysiology

A plastic head holder and three recording chambers were fixed to the skull under general anesthesia and sterile surgical conditions. Two of the recoding chambers were placed over the frontoparietal lobes in both hemispheres, tilted laterally by 35 deg, and aimed at the substantia nigra pars compacta (SNc) and the ventral tegmental area (VTA). The other recording chamber was placed over the midline of the frontal lobes, and aimed at the OFC and VS in both hemispheres. The head holder and recording chambers were embedded in dental acrylic that covered the top of the skull and were firmly anchored to the skull by plastic screws. After the surgery, the monkeys underwent a magnetic resonance imaging (MRI) scan to determine the position of recording electrode.

Single-unit recordings were performed using tungsten electrodes with impedances of 1.2 to 2.5 MΩ (Frederick Haer, ME, USA) that were introduced into the brain through a stainless-steel guide tube by an oil-driven micromanipulator (MO-97-S, Narishige, Tokyo, Japan). Recording sites were determined using a grid system, which allowed recordings at every 1 mm between penetrations. For a finer mapping of neurons, we also used a complementary grid that allowed electrode penetrations between the holes of the original grid. Based on obtained MRI images and using the grid system, we inserted the tungsten electrodes into the OFC, VS, and SNc/VTA (fig. S2). The OFC region of interest corresponded to area 13m, which ranged from A33 mm to A35 mm in monkey A and from A33 mm to A34 mm in monkey E along the anterior-posterior axis. The VS recording ranged from A23 mm to A30 mm in monkey A and from A22 mm to A28 mm in monkey E along the anterior-posterior axis.

Single-unit potentials were amplified and band-pass filtered (100Hz to 8 kHz) using a multichannel processor (MCP Plus 8, Alpha Omega, Nazareth, Israel) and isolated online using a voltage-time window discrimination system (ASD, Alpha Omega, Nazareth, Israel). The time of occurrence of each action potential was stored with 1-ms resolution.

### Identification of DA neurons

Putative DA neurons were identified based on their well-established electrophysiological signatures: a low background firing rate at around 5 Hz, a broad spike potential in clear contrast to neighboring neurons with a high background firing rate in the substantia nigra pars reticulata and a phasic excitation in response to free reward (*45, 46*).

### Statistical Analysis

For null hypothesis testing, 95% confidence intervals (*P* < 0.05) were used to define statistical significance in all analyses.

To examine the effects of actual and counterfactual outcomes on monkey’s next choice, we used a logistic regression analysis (Fig. 1D). We examined the effects separately for trials in which the monkey chose the first option (first option chosen trials) and trials in which the monkey did not choose it (first option unchosen trials). For each trial type, a logistic regression model was used to predict whether the monkey chose the first option in trial t+1 based on (1) the value of the first option in trial t+1, (2) the value of the first option in trial t, and (3) the value of the second option in trial t. The values of the first and second options in trial t are the values of the actual and counterfactual outcomes, respectively, in the first option chosen trials and vice versa in the first option unchosen trials. The models are expressed by the following equations:

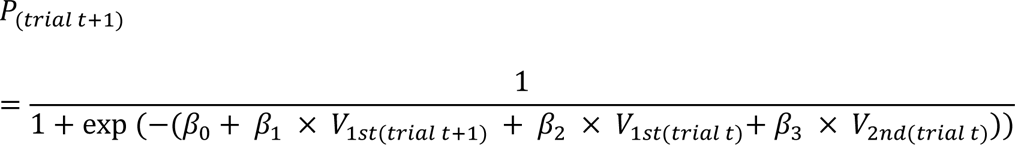

where P indicates the choice rate of the first option, *V_1st_* and *V_2nd_* indicate the value of the first and second options (1 to 6), and *β_0_*, *β_1_*, *β_2_*, and *β_3_* indicate the coefficients determined by logistic regression. Thus, *V_1st_* and *V_2nd_* in trial t are the values of the actual and counterfactual outcomes, respectively, in the first option chosen trials and vice versa in the first option unchosen trials.

To analyze neuronal activity, we combined the data obtained from the two monkeys because they were qualitatively identical.

To calculate spike density functions (SDFs), each spike was replaced by a Gaussian curve (σ = 30 ms).

To statistically identify the representation of counterfactual outcome value in OFC, VS and DA neurons, we used a calculation time window set after the second option onset during which neuronal modulation evoked by the second option was observed as population (DA: 100 – 400 ms; OFC and VS: 200 – 800 ms relative to the second option onset), and fitted the activity of each neuron during this window with a generalized linear model (GLM) separately for two decision trial types (first option chosen and unchosen trials) (Fig. 3 to 5). Given that the neuronal activity was also affected by the first option value, we put both the first and second option values into the model as two regressors. The model is expressed by the following equation:

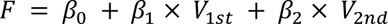

where *F* indicates the firing rate of each neuron, V_1st_ and V_2nd_ indicate the first and second option values (1 to 6), respectively, and *β_0_*, *β_1_* and *β_2_* indicate the coefficients determined by linear regression. If the fitted model contained a significant regression coefficient of the second option value in the first option chosen and unchosen trials, that neuron was considered to represent the actual and counterfactual values of the second option, respectively.

We used a bootstrap procedure to test whether the proportion of neurons showing a significant regression coefficient of the actual and counterfactual values of the second option was larger than expected by chance (Fig. 3A). For each neuron, the firing rate of each trial was shuffled and assigned to another trial at random to form a shuffled dataset. We fitted the firing rate of the shuffled dataset with the GLM and calculated the proportion of neurons showing a significant regression coefficient. We repeated such shuffling and calculation 1000 times and obtained the chance-level distribution of the proportion. We compared the original proportion of neurons showing a significant regression coefficient with the distribution.

We also used the GLM fitting method described above to calculate the temporal change in the proportion of neurons representing actual and counterfactual value of second option (Fig. 5A and C). Here the firing rate was calculated using a 100-ms sliding window with a 1-ms step, which was from −400 ms to 800 ms relative to the first and second option onsets. For each neuron, we defined the latencies of actual and counterfactual value signals as the start point of the first 50 consecutive 1-ms steps of which at least 45 steps exhibited a significant regression coefficient of each value (Fig. 3C).

## Acknowledgments

We thank the members of Naoshige Uchida laboratory for valuable discussions, E.S. Bromberg-Martin, I.E. Monosov, and L. Kingsbury for comments on an earlier version of the manuscript, and K. Bunzui for animal care.

## Author contributions

Conceptualization: MY, MM; Investigation: MY, MN, TK; Supervision: MM; Writing— original draft: MY, MM; Writing—review & editing: MY, MN, TK, JK, HY, HRK, MM

## Data and materials availability

All data needed to evaluate the conclusions in the paper are present in the paper and/or the Supplementary Materials. Additional data related to this paper may be requested from the authors.

