## Supplemental Information for "Distinct roles of the orbitofrontal cortex, ventral striatum, and dopamine neurons in counterfactual thinking of decision outcomes"

**The PDF file includes:**

Figs. S1 to S4

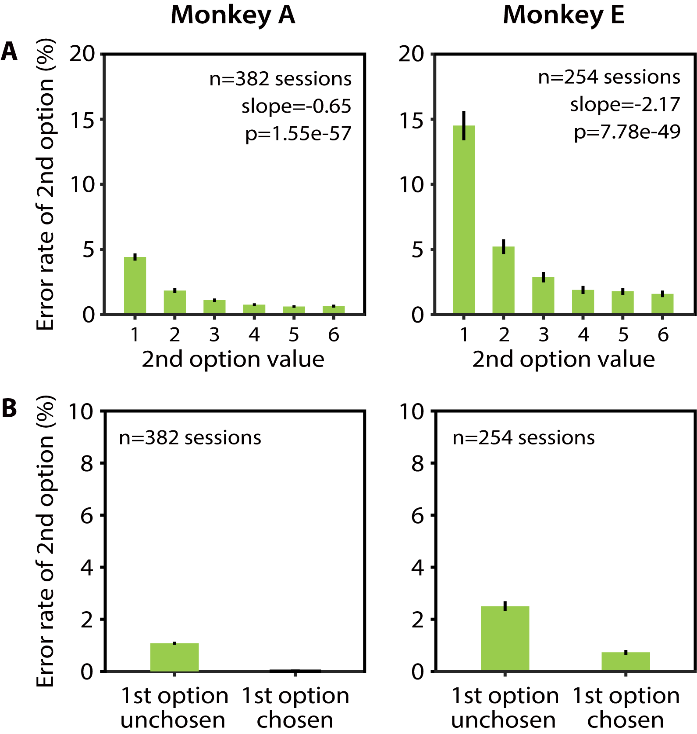

Fig. S1. Error rate during second option presentation. (A), Error rate during the presentation of the second option in the first option unchosen trials. Errors included failure to release the button, a break in central fixation, pressing the button again after releasing it. Data are shown as a function of the second option value in monkey A (left) and monkey E (right). Error bars indicate SEM. (B), Error rate during the presentation of the second option for the first option chosen versus unchosen trials.

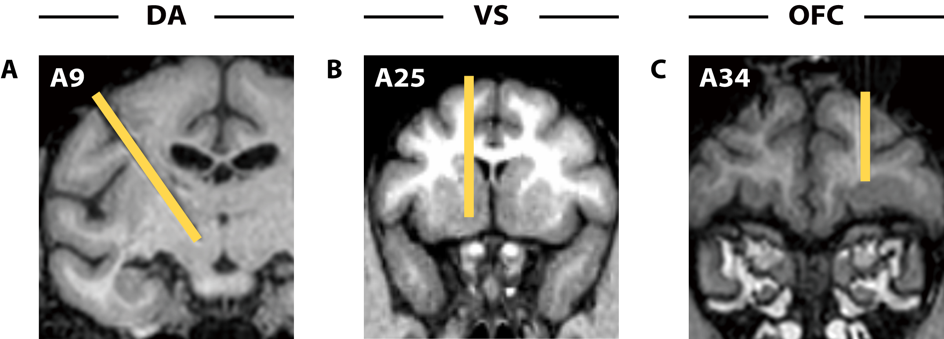

Fig. S2. Recording sites. (A to C), Representative penetrations of recording electrodes, shown via magnetic resonance imaging (MRI). Yellow bars indicate the electrodes targeting the left substantia nigra pars compacta/ventral tegmental area (SNc/VTA) (A) and the right orbitofrontal cortex (OFC) (C) in monkey E, and the left ventral striatum (VS) in monkey A (B).

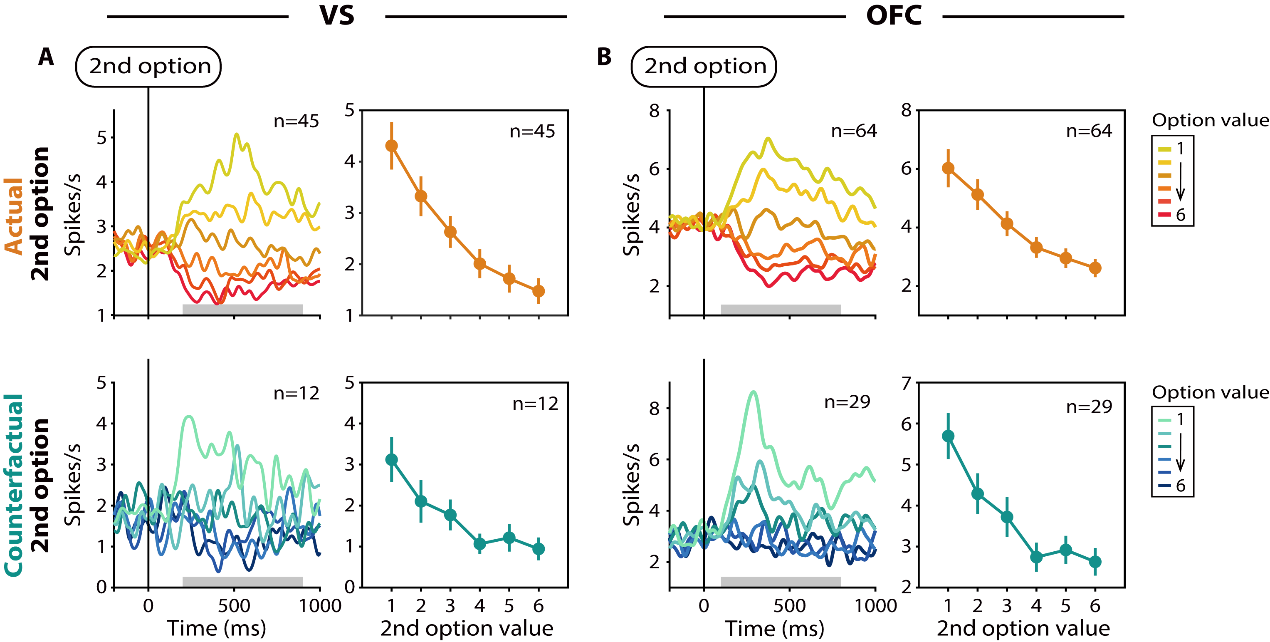

Fig. S3. Activity of VS and OFC neurons that negatively represented actual and counterfactual second option values. (A and B), Averaged SDFs aligned at the second option onset (left) and average magnitudes of neuronal activity evoked by the second option (right) for the first option unchosen and chosen trials in which the second option was the actual (top) and counterfactual (bottom) outcome, respectively. VS (A) and OFC (B) neurons that negatively represented the second option value were used in this analysis. Conventions are as Fig. 3B.

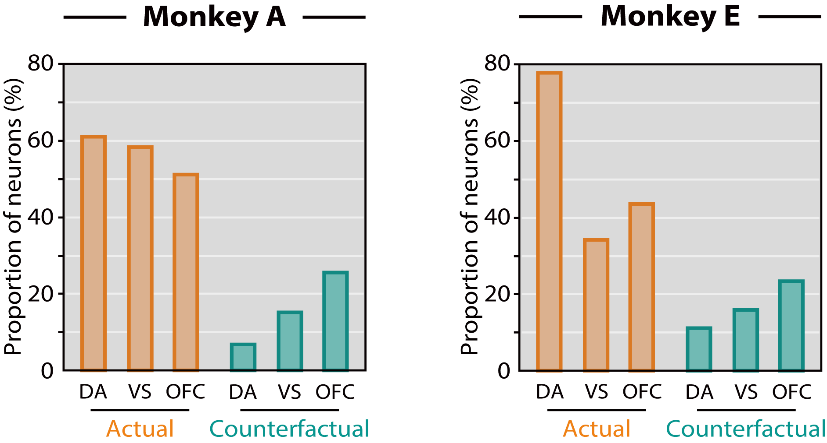

Fig. S4. Proportions of neurons representing the actual and counterfactual second option values in individual monkeys. Proportions of OFC, VS, and DA neurons representing the actual (orange) and counterfactual (green) values of the second option, shown separately for monkey A (left) and monkey E (right). The proportion of neurons representing the counterfactual value was not significantly different between the two monkeys (DA, *P* = 0.47; VS, *P* = 1.0; OFC, *P* = 0.77; Fisher’s exact test).
